# Spectral and non-spectral EEG measures in the prediction of working memory task performance and psychopathology

**DOI:** 10.64898/2026.03.25.714248

**Authors:** Fleming C. Peck, Catherine R. Walsh, Holly Truong, Jean-Baptiste F. Pochon, Kristen D. Enriquez, Carrie E. Bearden, Sandra K. Loo, Robert M. Bilder, Agatha Lenartowicz, Jesse Rissman

## Abstract

Working memory (WM) supports the temporary maintenance of goal-relevant information and is disrupted across many neuropsychiatric disorders. We examined whether scalp electroencephalography (EEG) data features beyond spectral power, including waveform shape, broadband spectral structure, and signal complexity, provide complementary information for predicting cognitive and clinical outcomes. EEG was recorded from 200 adults spanning a broad range of neuropsychiatric symptom severity while they completed three WM task paradigms: Sternberg spatial WM (SWM), delayed face recognition (DFR), and dot pattern expectancy (DPX). Separate machine learning models were trained on EEG features from the encoding, delay, and probe phase of each task to predict participants’ task accuracy, reaction time (RT) variability, WM capacity, and psychopathology scores (Brief Psychiatric Rating Scale). A split-half analytic framework was used, with cross-validated model development in an exploratory dataset (N=100) and evaluation of statistically significant models in a held-out validation dataset (N=100). In the exploratory dataset, SWM task data best predicted WM capacity, DPX task data predicted RT variability, and DFR task data predicted psychopathology, suggesting that these three WM paradigms engage distinct neural processes relevant to different outcomes. No models reliably predicted task accuracy. Models incorporating features beyond spectral power generally outperformed power-only models, and task-derived features outperformed resting-state-derived features. However, only those models predicting WM capacity and RT variability generalized to the validation dataset; models predicting psychopathology did not. These findings demonstrate functional heterogeneity across WM paradigms, show that complementary EEG features enhance predictive modeling, and highlight the importance of rigorous validation for identifying robust brain-behavior relationships.

## Introduction

Working memory is the cognitive capacity to temporarily store and manipulate goal-relevant information and plays a fundamental role in reasoning, problem-solving, and decision-making, forming the foundation for higher-order cognition (Baddeley, 1992). Successful working memory requires a balance between stable maintenance of information and flexible updating in response to changing environmental demands (Eriksson et al., 2015; Miller & Cohen, 2001; Myers et al., 2017). During a typical trial of a delayed response working memory task, stimuli must first be perceptually processed into task-relevant representations (encoding stage), then maintained in an active state for a short period of time (delay stage), and finally accessed and compared to a probe stimulus to facilitate a behavioral response (probe stage). These component processes impose different demands on neural systems, requiring coordination over time to support accurate working memory performance (D’Esposito & Postle, 2015).

Deficits in working memory are a hallmark of many neuropsychiatric disorders, including schizophrenia (Barch & Ceaser, 2012), attention-deficit/hyperactivity disorder (ADHD) (Martinussen et al., 2005), and depression (Snyder, 2013). Individuals with these conditions often struggle with maintaining and manipulating information in mind, leading to impairments in planning, attention, and executive functioning (McCabe et al., 2010). Importantly, working memory and sustained attention are tightly linked processes: effective working memory performance depends on the ability to maintain goal-relevant focus and resist lapses of attention over time (Chun, 2011; Esterman et al., 2013). Fluctuations in sustained attention, often indexed by reaction time variability, predict errors in working memory tasks and may represent a shared underlying control mechanism (Adam et al., 2015; Unsworth & Robison, 2018). These cognitive deficits frequently transcend traditional diagnostic categories, presenting instead as transdiagnostic symptom dimensions of varying severity (Bilder et al., 2013; McTeague et al., 2016).

Electroencephalography (EEG) provides a powerful tool for studying the neural basis of working memory, offering millisecond-level temporal resolution to track dynamic brain activity and clarify the relative contribution of different processes to working memory performance. Traditional EEG analyses have predominantly focused on oscillatory power – the amount of coordinated neural activity within specific frequency bands (e.g., theta, alpha, beta) – which has been central to many insights into working memory dynamics (for a review, see (Roux & Uhlhaas, 2014)). At the same time, power-based approaches make simplifying assumptions about signal stationarity and waveform sinusoidality. However, EEG signals reflect activity from multiple brain regions simultaneously (Klonowski, 2009), and neural oscillations often deviate from ideal sinusoidal patterns in ways that may carry meaningful physiological information (Cole & Voytek, 2017; Jones, 2016). These features of the signal, together with the inherently nonlinear and complex nature of brain dynamics (Bosl et al., 2011), present an opportunity to extract additional information that is not fully captured by power alone. Incorporating nonlinear and aperiodic measures of EEG signal offers a complementary framework for characterizing the temporal complexity, irregularity, and other dynamic properties underlying the working memory process.

For example, deviations from symmetric oscillations may reflect imbalances in excitatory and inhibitory processes, providing insight into circuit-level dynamics that support information processing (Barak & Tsodyks, 2014; S. R. Cole & Voytek, 2017; Kohonen, 2012). Aperiodic spectral features that quantify broadband, non-oscillatory components of neural activity may reflect global properties of neural excitability and network organization (Gao et al., 2017) and help to dissociate narrowband oscillatory effects from task-related shifts in the overall power spectrum (Donoghue et al., 2020). Additionally, measures of neural complexity capture the temporal diversity and irregularity of neural activity over time, providing a complementary index of the balance between stable and flexible neural dynamics (Lempel & Ziv, 1976). These measures have been highlighted as promising directions for memory research in recent reviews (Pavlov & Kotchoubey, 2022), with only a small number of recent applications (e.g., (Bender et al., 2025; Frelih et al., 2025; McKeon et al., 2024; Virtue-Griffiths et al., 2025)). A key goal of the present project is to establish their physiological significance by evaluating their relationship to working memory performance, thereby advancing our understanding of the neural mechanisms that support working memory function.

Meta-analytic evidence indicates that alterations in EEG power are rarely disorder-specific but instead reflect shared disruptions in neural regulation across neuropsychiatric diagnoses, as well as substantial variability within diagnoses (Newson & Thiagarajan, 2018). Consistent with this transdiagnostic perspective, we examined a heterogenous sample spanning a range of clinical diagnoses and symptom severity, allowing brain-behavior relationships to be assessed along a continuous spectrum rather than within categorical diagnostic boundaries. From a methodological standpoint, brain-based prediction benefits from broad variance in both neural and behavioral measures. Accordingly, a mixed sample expands the range of outcomes and increase sensitivity for modeling individual differences in working memory ability and related cognitive processes.

The present investigation integrates measures of complexity, oscillation shape, and aperiodic spectral components with the standard power analyses to provide a detailed characterization of the EEG-based neural dynamics supporting working memory. To relate these neural features to behavior, we use ridge regression, a regularized machine learning model that reduces instability due to collinearity among correlated predictors, with cross-validation to promote generalization to new data (Hoerl & Kennard, 1970). Models were trained to predict a series of behavioral outcomes, from working memory task-related measures of task accuracy or sensitivity and reaction time variability to more distal measures indexing cognitive and clinical traits derived from neuropsychological testing. Analyses were conducted in a large, clinically heterogenous sample using a cross-validated exploratory design in one half of the data and independent validation in the other half to assess model generalizability. This approach allowed us to evaluate whether an expanded set of EEG features provide complementary predictive information to power alone, to assess how distinct working memory task paradigms relate differently to behavioral outcomes, and to compare the predictive utility of task-evoked versus resting-state EEG in linking brain activity to cognition and psychopathology.

## Methods

### Participants

A total of 200 adults were recruited as part of the “Multi-Level Assays of Working Memory and Psychopathology” study supported by the National Institute of Mental Health Research Domain Criteria (RDoC) Initiative (R01-MH101478). The sample included 150 care-seeking individuals and 50 non-care-seeking individuals to ensure a broad range of psychopathology and symptom variability. EEG data from this sample have been previously analyzed using standard analyses of oscillatory power and ERPs in separate reports (Lenartowicz et al., 2019, 2021). The care-seeking group included individuals seeking treatment for mental or emotional health concerns, recruited through advertisements or referrals from UCLA’s Neuropsychiatric Behavioral Health Service. The non-care-seeking group consisted of individuals who had not sought behavioral health or substance-use treatment within the past year. All participants completed the same behavioral and clinical assessments and were analyzed together to increase variance in cognitive performance and symptom severity, enabling dimensional analyses of brain-behavior relationships.

Inclusion criteria required participants to be: (1) between the ages of 21 and 40, (2) have at least eight years of formal education, (3) be proficient in English, (4) have an IQ above 70 (assessed via the WAIS-IV Vocabulary and Matrix Reasoning subtests), and (5) possess normal vision (20/50 or better in each eye). Exclusion criteria included: (1) a history of medical or neurological illness or treatment known to affect cognition, (2) use of psychotropic or sedating drugs within 24 hours prior to testing, (3) recent administration of long-acting antipsychotics or electroconvulsive therapy (within the past six months), (4) a diagnosis of a substance use disorder (excluding caffeine or nicotine) within the past six months, (5) positive urinalysis for THC, cocaine, amphetamines, opiates, or benzodiazepines, and (6) contraindications for MRI scanning. All participants completed comprehensive diagnostic and neurocognitive assessments as part of the study. Demographic characteristics of the sample, including age and sex, are reported below as a function of the split-half exploratory and validation datasets (see *Split-half dataset characteristics*)

### Tasks

EEG data were recorded while participants completed four working memory tasks: Sternberg spatial working memory (SWM), delayed face recognition (DFR), dot pattern expectancy task (DPX), and lateralized change detection (LCD). Resting-state data were also collected. The tasks were presented in two orders, counterbalanced across participants: SWM, LCD, eyes-closed rest, DPX, eyes-open rest, DFR or DFR, eyes-closed rest, DPX, eyes-open rest, LCD, SWM. LCD was excluded from the present study because trial duration was determined to be insufficient to get reliable neural measure estimates of each working memory sub-process. Example trials from each included task are visualized in Figure 1. All tasks were controlled by E-Prime 3.0 software (Psychology Software Tools, Inc., 2016) run on a Dell PC, and stimuli were presented on a 17” monitor with responses collected on a keyboard.

**Figure 1:**
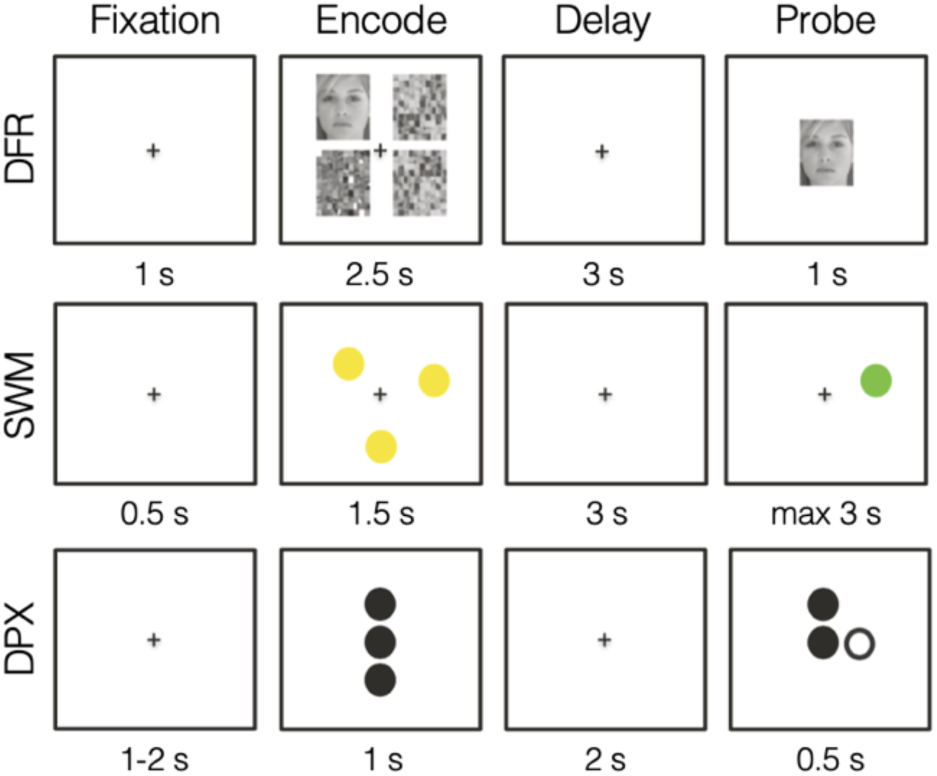
Trial structure and stage duration for working memory tasks. Example trials are shown for the three working memory tasks: Delayed Face Recognition (DFR; top), Spatial Working Memory (SWM; middle), and Dot-Probe Expectancy (DPX; bottom). All tasks comprise the same sequence of sub-stages: Fixation, Encode, Delay, and Probe. The duration of each stage is indicated beneath the corresponding visual. The top panel illustrates a low-load DFR trial with a matching probe, the middle panel shows a load=3 SWM trial with a matching probe, and the bottom panel depicts an AX trial in the DPX task.

The SWM task is a delayed match-to-sample adaptation of the Sternberg paradigm (Sternberg, 1966), a well-established test of working memory maintenance. Trials consist of 0.5s of initial fixation, a 1.5s screen with yellow dots of which participants are tasked with remembering their locations, a 3s blank screen requiring participants to maintain dot location(s) in memory, and a 3s probe screen with a single yellow dot for which participants indicate if this probe stimulus matches any of the dots shown previously in the trial (match) or not (non-match). Participants indicate their choice with a button press, which triggers the end of the trial, followed by a 1.5-2s intertrial interval. This paradigm included four load conditions: 1, 3, 5, and 7 yellow dots to be encoded. Participants completed 24 trials per load (96 total) with these trials evenly distributed across four blocks of 24 trials each.

The DFR task is another delayed match-to-sample task but with face stimuli as the memoranda. Trials consist of low load (one face) or high load (three faces) shown for 3s with scrambled faces in the unoccupied quadrants, a 3s delay period, a 1s probe face where participants respond with a button press whether it matches any of the faces encountered on that trial, followed by 1s intertrial interval. The gender of the faces (either all male or all female within a given trial), memory load (low or high), and match or nonmatch probe conditions were balanced across the experiment (50% of each type), and trials of these varying conditions occurred in a randomly intermixed order. Participants performed a total of 96 trials, presented in three blocks of 32 trials.

The DPX task is a continuous performance task variant of the AX-CPT task used to assess working memory goal maintenance and developed as a clinical tool (J. D. Cohen et al., 1999). The adaptation translates the original letter stimuli to dots and reduces the trial counts to minimize testing time while maintaining sensitivity. The task consists of two contexts, A and B, and two stimuli, X and Y. Trials in the task require participants to keep track of both a context and a stimulus and modulate their response in consideration of the conjunction of the two variables. Specifically, participants were to respond with a button press only for AX trials and not at all for AY, BX, or BY trials. A trial consisted of a 1s presentation of the context stimulus, a 2s maintenance interval, a 0.5s presentation of the target stimulus, and a 1-2s intertrial interval. Participants were therefore required to maintain the context to inform their target response after a short delay. Participants completed a total of 216 trials: 156 AX trials, 24 AY trials, 24 BX trials, and 12 BY trials. These trial types were evenly distributed over six blocks of 36 trials each.

Eyes-open (EO) and eyes-closed (EC) resting-state EEG data were recorded. Five minutes of data were collected for each condition. Participants were instructed to sit still and relax throughout both recordings. In the EO condition, a screensaver video consisting of moving circles was played. In the EC condition, participants were asked to close their eyes and to try not to fall asleep.

### EEG recording

EEG data were recorded using a BioSemi ActiveTwo system with 64 Ag/AgCl electrodes positioned according to the 10-20 system. Electrode impedances were maintained below 10 kΩ before task initiation. Signals were acquired in a DC-coupled mode at a sampling rate of 1024 Hz with no online referencing using BioSemi hardware and ActiView recording software (BioSemi B.V., Netherlands).

### EEG preprocessing

EEG data were preprocessed in MATLAB (MathWorks, Inc., Natick, MA) using functions from EEGLAB (v11.03.b). Signals were high-pass filtered at 1 Hz, and visually identified noisy channels were excluded from further analysis. Data were then re-referenced to the average reference. Task EEG signals were epoched into individual trials, while resting-state data were segmented into 2-second epochs. Within each participant, epochs were excluded from analysis if signal power exceeded the 85^th^ percentile across more than 60% of channels, as these were considered to reflect gross movement artifacts or muscle activity.

### EEG feature extraction

EEG measures were computed using data from 12 electrodes: F3, Fz, F4, C3, Cz, C4, P3, Pz, P4, O1, Oz, and O2. The EEG data were segmented in trial epochs and filtered into canonical frequency bands: theta (3–7.9 Hz), alpha (8–11.9 Hz), beta (12–29.9 Hz), and gamma (30–55 Hz). Trial epochs were then further segmented by task stage (encoding, delay, probe) to ensure neural activity was analyzed in relation to specific cognitive processes. Filtering was applied prior to stage segmentation to minimize edge distortions or artificial signal amplification. To maximize the number of trials contributing to the estimation of EEG signal measures, data from all trials were included and collapsed across task condition (e.g., high vs. low load; matching vs. non-matching probe) and behavioral accuracy. A summary of the EEG features included in the study is included in Table 1, and more detailed descriptions of computation are described below.

**Table 1:** EEG measures with qualitative description.

| **Construct** | **Measure** | **Description** |
| --- | --- | --- |
| Spectral power | Normalized power | Quantifies the magnitude of sinusoidal neural activity relative to baseline activity levels. |
| Waveform symmetry | Peak-trough symmetry;<br>Rise-decay symmetry | Quantifies asymmetries in oscillatory waveforms reflecting non-sinusoidal cycle shape. |
| Oscillation monotonicity | Monotonicity | Quantifies the smoothness and consistency of rise and decay phases within oscillatory cycles. |
| Cycle consistency | Amplitude consistency;<br>Period consistency | Quantifies the stability of oscillatory strength and timing across successive cycles. |
| Power spectrum structure | Spectral exponent;<br>Spectral offset | Quantifies the distribution of frequency power within the power spectrum. |
| Complexity | Lempel-Ziv complexity | Quantifies temporal irregularity and pattern diversity over time. |

Several properties of individual neural oscillations were computed using the open-source ByCycle package (S. Cole & Voytek, 2019). Monotonicity measures the proportion of voltage changes that are positive during the rise phase and negative during the decay phase. Oscillation symmetry was assessed by examining two key metrics: rise-decay symmetry, calculated as the ratio of rise duration to the total oscillation period, and peak-trough symmetry, defined as the ratio of trough duration to the total period. To capture overall symmetry, we computed the absolute difference from 0.5, where 0.5 represents perfect symmetry. Amplitude consistency was calculated as the ratio of rise and decay voltages, with values deviating further from 1 indicating greater inconsistency. Period consistency was defined as the maximum deviation in oscillation period compared to preceding and following cycles. These measures were computed for each oscillation and then averaged within each EEG segment.

Lempel-Ziv Complexity (LZC; (Lempel & Ziv, 1976)) was used to assess the randomness and structure of neural signals. First, the EEG signal was binarized using the median voltage as a threshold, ensuring an equal distribution of 0s and 1s. Unique patterns were then identified by scanning forward to detect the shortest substring not previously encountered. The total count of unique patterns determined the raw LZC score, with higher values indicating greater signal variability. To control for differences in sequence length, the raw LZC score was normalized by dividing it by the expected complexity of a random sequence of the same length.

A Fourier transform was applied to the unfiltered EEG signal to compute the power spectrum. Power estimates were obtained for each frequency band by averaging the power of oscillations within the respective range. For task-related epochs, power was normalized using the mean power 900–300 ms before encoding onset as a baseline to maintain consistency with prior task-based EEG studies. Power estimates from the resting-state sessions remained unnormalized.

We used the open-source Spectral Parameterization package (formerly FOOOF) to characterize broadband shape of the power spectrum (Donoghue et al., 2020). The power spectrum was log-transformed, and its aperiodic component was modeled using a power-law function of the 1/f distribution. The spectral offset reflects global shifts in power, while the spectral exponent quantifies the steepness of spectral decay. Evaluating these global power spectrum components helps prevent misinterpretation of local power fluctuations that may stem from broader spectral changes. Unlike oscillatory power estimates, which were calculated separately for each frequency band, aperiodic features were derived across the full 3–55 Hz range.

To enhance signal reliability and reduce noise, features were first averaged across all trials within each channel and task stage. These channel-level values were then aggregated into four electrode clusters, with each defined as the mean of three electrodes spanning the scalp: frontal (F3, Fz, F4), central (C3, Cz, C4), parietal (P3, Pz, P4), and occipital (O1, Oz, O2).

### Behavioral outcome measures

Models were trained to predict outcome measures spanning three key domains: task-related performance, working memory capacity, and neuropsychiatric symptomatology. Pairwise Pearson correlations among all behavioral outcome measures are provided in Supplemental Figure S1A.

To quantify task-specific working memory performance, we derived two behavioral metrics from each paradigm: accuracy and reaction time (RT) variability. Accuracy reflected participants’ ability to encode, maintain, and retrieve information and was operationalized as overall accuracy (proportion correct across all conditions) for the SWM and DFR tasks and as sensitivity (d-prime; z transformed hit rate minus z transformed false alarm rate) for the DPX task. RT variability was calculated as the standard deviation of RTs (ms) across all trials.

A composite measure of working memory capacity was obtained from standardized cognitive assessments. The score on six subtests was averaged. Four were included from the Wechsler Adult Intelligence Scale – Fourth Edition (WAIS-IV) subtests: Letter-Number Sequencing, which requires participants to reorganize sequences of letters and numbers in ascending and alphabetical order; Digit Span Forward, where participants repeat numbers in the presented order; Digit Span Backward, which requires reversing the sequence; and Digit Span Sequencing, which involves repeating numbers in ascending order. Two were included from the Wechsler Memory Scale – Fourth Edition (WMS-IV): Symbol Span, in which participants recreate a sequence of abstract symbols after a brief delay, and Spatial Addition, which requires mentally combining positions of dots presented separately in two grids. Each subtest score was converted to proportion correct, and the mean of these rescaled subtests was used as a composite index of working memory capacity.

To examine more specific neuropsychiatric symptomatology, we include several self-reported measures of psychological distress and cognitive function. Symptoms of Anxiety and Depression were assessed using the Patient-Reported Outcomes Measurement Information System (PROMIS; Cella et al., 2010), a self-report survey that measures emotional distress over the past seven days on a 5-point Likert scale. Standardized T-scores that represent severity relative to the general population were included. Broader psychopathology was evaluated using the Brief Psychiatric Rating Scale (BPRS; Overall & Gorham, 1962). Four BPRS factor scores were computed using factor structures identified in a large-scale meta-analysis of prior BPRS factor analytic studies (Dazzi et al., 2016): Positive Symptoms, Negative Symptoms, Activation, and Affect. Factor scores for each participant were calculated by aggregating item responses according to this published factor solution. Because substantial floor effects (i.e., scores at or near the minimum of the scale) were observed for the Positive Symptoms (66.7% of participants), Negative Symptoms (66.7% of participants), and Activation (33.7% of participants) factors, only the Affect factor was included as a behavioral outcome. The Affect factor integrates across items pertaining to anxiety, guilt, depression, and suicidality and exhibited meaningful variability in the sample. The BPRS total score was also analyzed as a global index of psychopathology, representing the cumulative severity of symptoms across all domains and serving as a proxy for overall psychiatric burden within our clinically heterogeneous sample (Hofmann et al., 2022). Notably, we do not attempt to predict categorical psychiatric diagnoses (e.g., major depressive disorder) but instead focus on symptom severity as a continuous measure. This transdiagnostic dimensional approach allows for a more precise investigation of the relationships between brain activity and specific symptoms, avoiding the limitations of broad, heterogeneous diagnostic labels.

### Split-half dataset characteristics

To ensure the robustness and generalizability of our findings, we partitioned the N=200 sample into discovery and replication cohorts, where models were developed on one half of the data and subsequently tested on a strictly held-out validation set. The exploratory discovery dataset (N=100) was used to evaluate a broad range of models linking measures of EEG activity to behavioral outcomes, while the validation dataset (N=100) was reserved to confirm the generalizability of only the most promising models identified in the exploratory analyses. By freezing our models prior to attempting replication on a completely untouched dataset, we mitigate the risk of overfitting and the winner’s curse that plagues many efforts to predict behavior from neuroimaging data (Hosseini et al., 2020; Vul et al., 2009). This prospective validation approach thus enhances the credibility and external validity of our ultimate findings. The split was pseudo-randomized to preserve gender distributions within each diagnostic category: no clinical diagnosis (28 M, 30 F), major depressive disorder (26 M, 61 F), anxiety disorder (12 M, 26 F), schizophrenia (4 M, 4 F), anorexia (1 F), antisocial personality disorder (1 M), and ADHD (2 M, 5 F). Diagnoses reflect each participant’s primary clinical diagnosis, though some comorbidity was present in the sample. This procedure yielded an exploratory dataset with 36 males and 64 females (mean age of 27.8 ± 5.2 years) and a validation dataset with 37 males and 63 females (mean age of 28.3 ± 5.4 years).

A total of seven participants were excluded from EEG analyses. Five participants were removed due to poor signal quality associated with thick hair, which impeded electrode contact, and two participants were excluded for excessive drowsiness and inattention during the experimental sessions. Additionally, two participants had missing task data (one for DFR and one for SWM). The final exploratory dataset included 95 participants (94 for SWM), and the validation dataset included 98 participants (97 for DFR).

### Model development

Models based on task data consisted of 360 features, each defined by the combination of EEG measure, frequency band, electrode cluster, and task stage. Specifically, seven EEG measures were computed across four frequency bands (amplitude consistency, period consistency, rise-decay symmetry, peak-trough symmetry, monotonicity, power, complexity; 28 features) and two across the full spectrum (spectral exponent and offset; 2 features), averaged within four electrode clusters (30 x 4 = 120 features), and computed for each of the three task stages (120 x 3 = 360 features). This is visualized in Figure 2. Resting-state models included only 120 features, corresponding to a single stage (e.g., rest).

**Figure 2:**
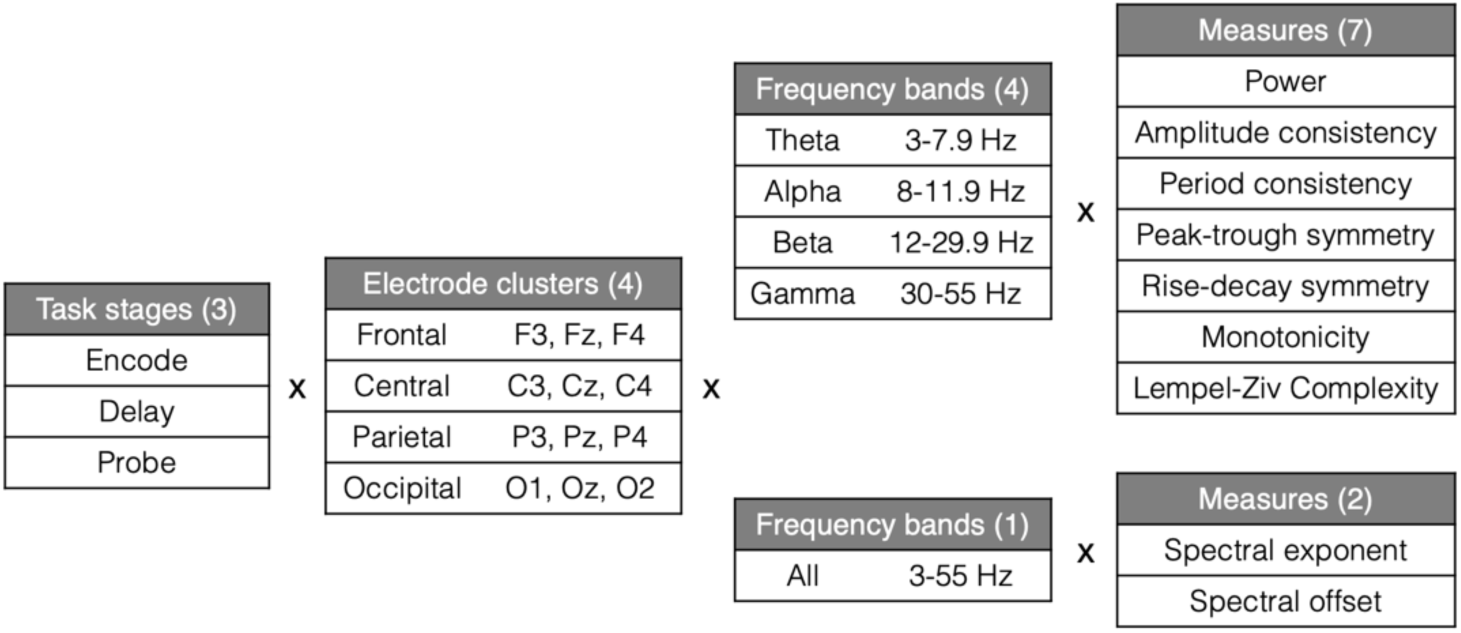
EEG feature dimensions. Overview of the feature dimensions of model inputs. Each individual feature corresponds to a specific EEG measure computed from signal filtered to a given frequency band, averaged within an electrode cluster, and calculated separately for each task stage.

Ridge regression was selected for model development because it performs robustly in high-dimensional, multicollinear feature spaces typical of EEG data. Models were implemented in Python using the RidgeCV function from the scikit-learn library (Pedregosa et al., 2018). This approach performs ridge regression with internal cross-validation to select the optimal regularization parameter (α). Ridge regression consistently yielded the highest or comparable predictive accuracy across tasks and behavioral outcomes relative to alternative models, including LASSO regression, ordinary least squares, and linear support vector machine. Performance for the outcome variables selected for validation is shown in Supplemental Figure S3. Based on this comparative performance, ridge regression was used for subsequent analyses.

Exploratory models were trained using 10-fold cross-validation, a procedure designed to assess generalizability while reducing the risk of overfitting. The exploratory dataset was randomly divided into 10 equally sized folds. The model is trained on data from nine folds and then tested on the remaining fold, ensuring that no participant is ever used for both training and testing across different iterations. Prior to model fitting, features were z-scored using the mean and standard deviation computed from the training folds, and these parameters were then applied to standardize the test fold to prevent information leakage. This process is repeated 10 times, with each fold serving as the test set once, and overall performance is averaged across these iterations. Since cross-validation results can be influenced by how participants are assigned to folds, we repeat the entire cross-validation procedure 30 times with different randomized fold assignments.

For final evaluation of promising exploratory models, a single model was trained on the full exploratory dataset and then tested on the independent validation dataset.

### Model performance assessment

Model performance was quantified as the Spearman correlation between predicted and observed values across all participants (e.g., after concatenating the model’s predictions across all folds in the exploratory analyses). To obtain a stable overall estimate, correlation values were Fisher z-transformed prior to averaging across iterations and then inverse-transformed for reporting.

To determine whether model performance was statistically meaningful, a null distribution was generated by randomly shuffling outcome variables and testing how well the model predicted these permuted values 1,000 times. The observed model performance correlation was considered statistically significant if it exceeded 95% of the values in this null distribution, indicating that the model captured systematic relationships between neural features and behavioral outcomes rather than random noise.

### Validation performance assessment

For each behavioral outcome, the model demonstrating the strongest statistically significant performance in the exploratory analyses was selected for independent validation. For each selected outcome, a final model was trained using all participants in the exploratory dataset and then applied to the held-out validation dataset to generate predicted outcome values. Validation performance was quantified as the Spearman correlation between predicted and observed values across validation participants.

Statistical significance was assessed using a permutation-based null distribution generated by randomly shuffling outcome values within the validation dataset 1,000 times. Observed performance exceeding the 95th percentile of the null distribution was considered statistically significant, indicating successful generalization to independent data.

### Feature importance assessment

Feature importance was estimated from models trained on all exploratory dataset participants (e.g., without cross-validation) because the goal was to examine feature contributions rather than evaluate model performance. To establish a baseline, a null weight distribution was generated by permuting the outcome variables and retraining the model 1,000 times, extracting the feature weights from each iteration. Statistical significance was determined using two-tailed tests: for example, with alpha = 0.05, a feature was considered significant if its observed weight fell below the 2.5^th^ or above the 97.5^th^ percentile of this null distribution. Feature importance analyses were evaluated only in models that significantly generalized to the independent validation dataset.

## Results

We first examined relationships among EEG measures. Figure 3 shows the average pairwise correlation between EEG measures across electrode clusters, frequency bands, task stages, and tasks in the exploratory dataset. Correlations were computed between measures across these dimensions. Self-comparisons of identical measures (which would yield correlations of 1) were excluded. For each pair of measures, correlations were averaged across all combinations of electrode clusters, frequency bands, task stages within each task and then averaged across tasks. Each cell therefore reflects the average correlation between two measures computed across these feature dimensions. Diagonal values do not reflect self-correlations but instead reflect the average correlation between different combinations of electrode cluster, frequency band, task stage, and task for the same measure. Power showed modest association with itself (average r = 0.3 across these feature dimensions) and was largely uncorrelated with the other measures. This pattern supports our hypothesis that the additional signal processing metrics capture complementary aspects of neural activity beyond power modulations. Correlations among behavioral outcome measures and comparisons of outcome distributions between the exploratory and validation datasets are shown in Supplemental Figure S1.

**Figure 3.**
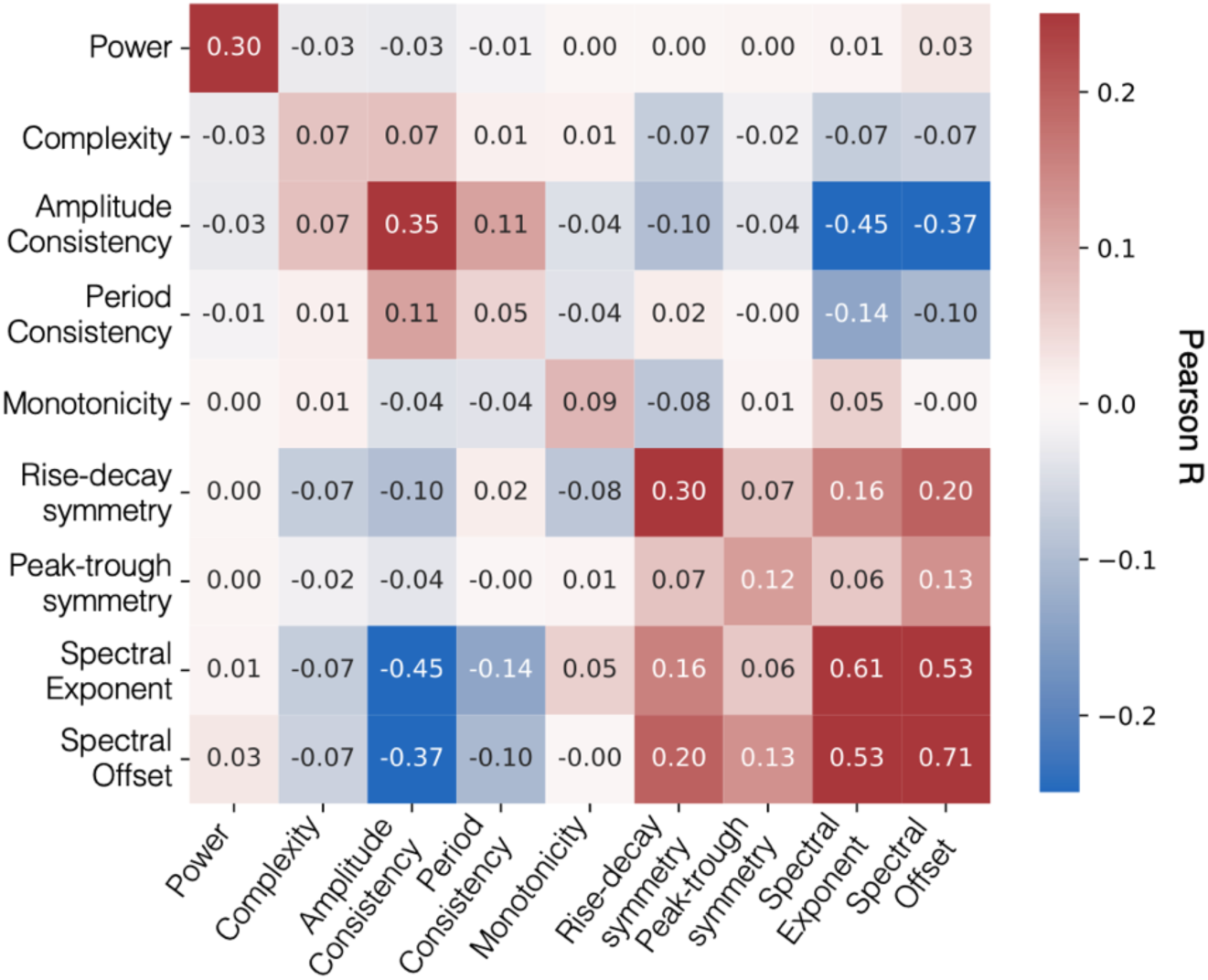
Correlations between EEG measures. Each cell of the correlation matrix shows the average Pearson correlation between two EEG measures, computed across electrode clusters, frequency ranges, and task stages. Correlation matrices were first generated separately for each task and then averaged across the three tasks. Cell values indicate Pearson’s *r*, and the color represents both magnitude and sign of the correlation.

Next, we evaluated the ability of cross-validated machine learning models to predict behavioral outcomes with our exploratory dataset half (n=100). Figure 4 presents modeling results examining how EEG features computed from each task stage predict behavioral outcomes. Behavioral measures varied in their proximity to the EEG task, ranging from directly-related measures derived from the very tasks that participants were performing as EEG data were collected (task accuracy, RT variability) to broader assessments of cognitive and clinical traits (WM capacity and psychopathology). Demographic control models including only age and gender as predictors were not significant for any behavioral outcome (all *p*’s > 0.05), so these variables were not regressed out or included in subsequent analyses. None of the models predicted task performance (accuracy or d-prime) for any of the tasks. However, EEG features from the DPX task significantly predicted RT variability on the DPX task (Spearman’s rho = 0.47; *p* < 0.001), and features from the SWM task significantly predicted WM capacity as measured by the behavioral tasks (Spearman’s rho = 0.32; *p* = 0.014). Overall psychopathology (BPRS total score) was significantly predicted by features from the DFR (Spearman’s rho = 0.40; *p* = 0.001) and DPX (Spearman’s rho = 0.25; *p* = 0.032) tasks, and Affect was significantly predicted by DPX features (Spearman’s rho = 0.32; *p* = 0.016). Subsequent analyses focus on the strongest predictive model for each behavioral outcome. Accordingly, for overall psychopathology, the DFR model showed the strongest predictive performance and was therefore pursued in follow up analyses despite significant prediction from more than one task.

**Figure 4.**
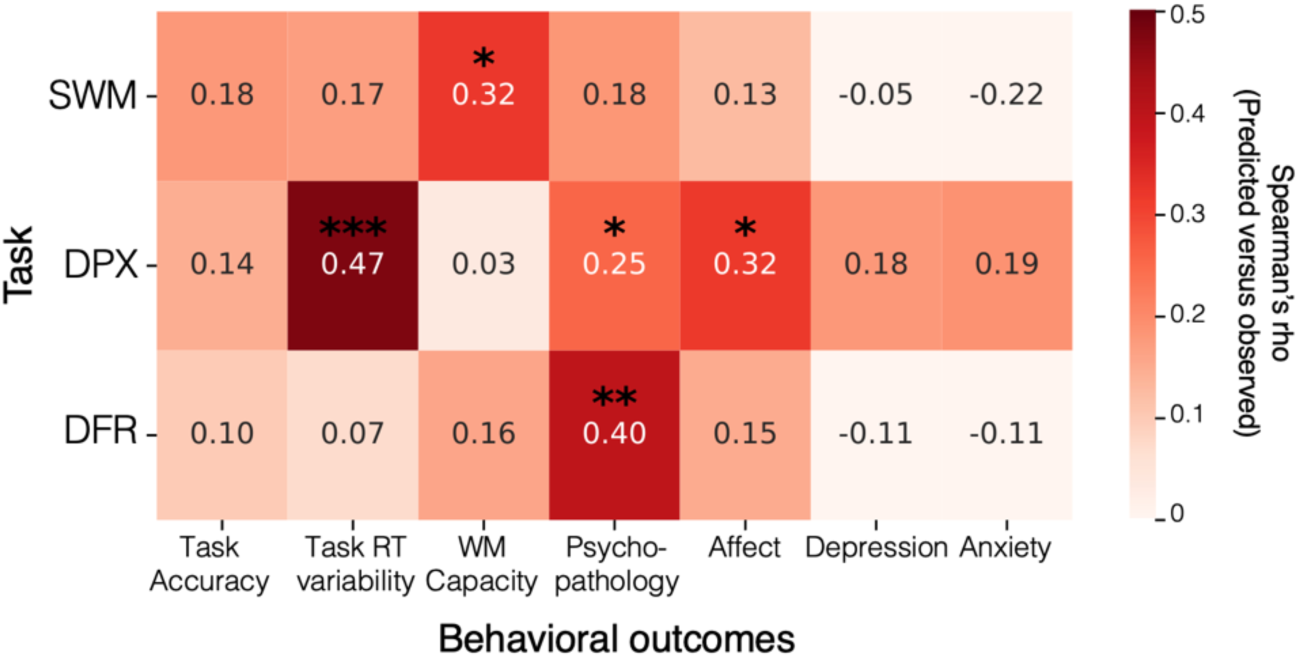
Exploratory model performance. Heatmap of the Spearman correlation between observed and predicted values for models trained with EEG data during a given task (y-axis) for behavioral outcome variables (x-axis). Color reflects strength of correlation that is also denoted by the value in the center of each cell. *p<0.05; **p<0.01; ***p<0.001 (relative to proportion greater than null permutation distribution).

The models reported thus far used EEG measures computed separately for each task stage (encoding, delay, and probe) and concatenated into a single feature space. To assess whether activity during individual stages could independently predict behavior, we also evaluated stage-specific models (Supplemental Figure S2). Predictive strength varied across stages, indicating that behaviorally relevant neural signals are not uniformly sustained throughout the trial but instead reflect process-specific dynamics tied to distinct cognitive operations. For the SWM task, none of the stage-specific models significantly predicted working memory capacity, suggesting that the predictive utility of the combined model arises from integrating information distributed across stages rather than from any single stage alone. These observations motivated our focus on models that integrate information across multiple task stages.

We next evaluated whether model performance could be explained by power alone versus models incorporating additional EEG features and resting-state characteristics. To this end, we predicted each behavioral outcome using models trained on only power, all measures except for power, and resting-state EEG features (Figure 5A). Across behavioral outcomes, models that included the additional measures (colored bars) consistently outperformed those based solely on power (dark gray bar), highlighting the value of incorporating richer aspects of neural dynamics. Indeed, in nearly all cases, the inclusion of all features together (colored bars) performed better than using all other features outside of power (light gray bars). The model predicting WM capacity from SWM features is the exception, where the model excluding power features showed numerically stronger prediction accuracy compared to when they were included. Models with power features alone remained significant for this task, albeit with reduced predictive strength. Finally, the psychopathology models were significantly predicted by eyes-open resting-state features (downward striped bars), suggesting that psychiatric symptomatology may reflect a relatively stable, trait-like characteristic rather than a transient, task-evoked state.

**Figure 5.**
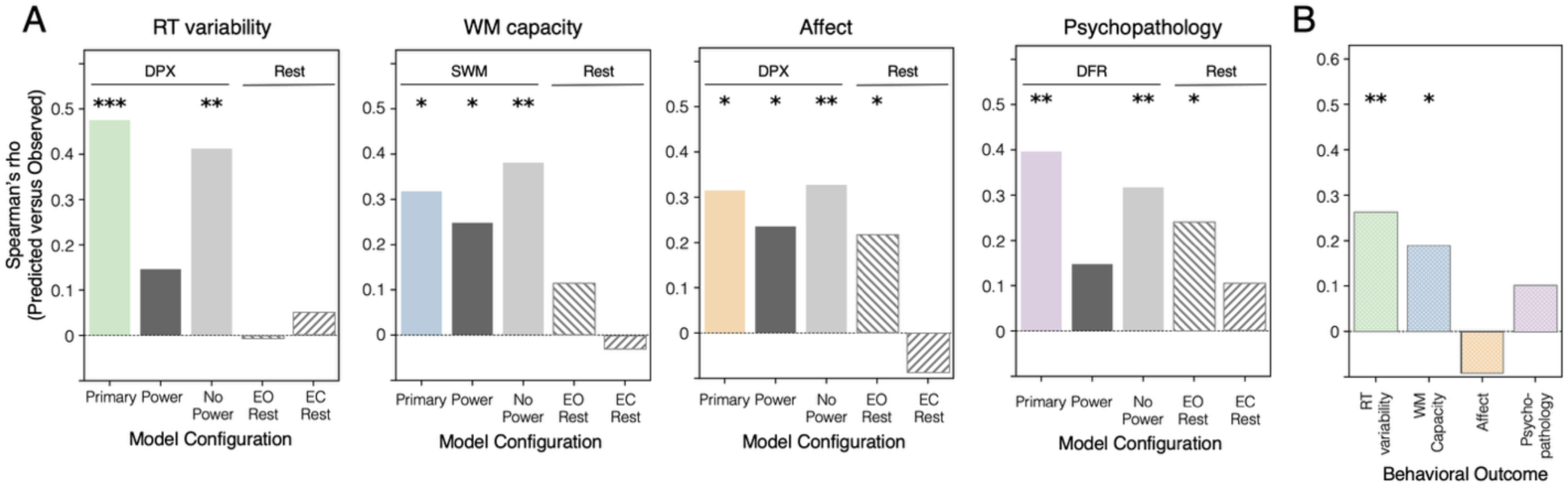
Validation of model generalization. Bar plots show model performance operationalized as Spearman correlation between observed and predicted values. **(A)** Model performance for model of interest using all features (“primary”) shown in colored bar for DPX data predicting reaction time variability, SWM data predicting WM capacity, DPX data predicting Affect, and DFR data predicting overall psychopathology. Four model comparisons are included for each analysis: same task data and outcome measure but with only power-derived features (dark gray, “power”); same task data and outcome measure but with all measures except for power (light gray, “no power”); same outcome measure predicted by all features derived from eyes-open (EO) resting-state data (downward right lines); and same outcome measure predicted by all measures derived from eyes-closed (EC) resting-state data (upward right lines). Horizontal annotations above plots indicate task-based and resting-state EEG models. **(B)** Validation model performance trained with exploratory dataset and tested on held-out dataset: DPX data predicting RT variability (green); SWM data predicting WM capacity (blue); DPX data predicting Affect (orange); and DFR data predicting psychopathology (purple). The models predicting Affect and overall psychopathology did not generalize to the validation dataset (*p* > 0.05). \**p*<0.05; \*\**p*<0.01; \*\*\**p*<0.001 (relative to proportion greater than null permutation distribution).

The primary goal of our analysis was to evaluate the generalizability of the strongest models using completely held-out validation data (Figure 5B). Of the four models that showed the most promising cross-validated performance within the exploratory dataset, two successfully generalized, such that they were able to reliably predict outcomes in the validation dataset. The two successful models were the ones predicting cognitive outcomes: DPX features predicting RT variability on the DPX task (Spearman’s rho = 0.26; *p* = 0.002) and SWM features predicting WM capacity (Spearman’s rho = 0.19; *p* = 0.027). In contrast, both models relating task EEG data to clinical symptomatology did not generalize to the validation dataset: the model predicting Affect showed no correspondence between predicted and observed scores (Spearman’s rho = −0.09, *p* = 0.82), and the DFR model predicting overall psychopathology was also nonsignificant (Spearman’s rho = 0.10; *p* = 0.14). Scatterplots depicting model fits for both the exploratory and validation datasets are provided in Supplemental Figure S4.

Finally, we examined which neural features contributed most strongly to the two models that successfully generalized across dataset halves in an effort to gain deeper understanding of which properties of neural activity account for individual differences in behavior (Figure 6). All features were z-scored prior to model fitting, so coefficient magnitudes are comparable within models but not across, given differences in behavioral outcome scales.

**Figure 6.**
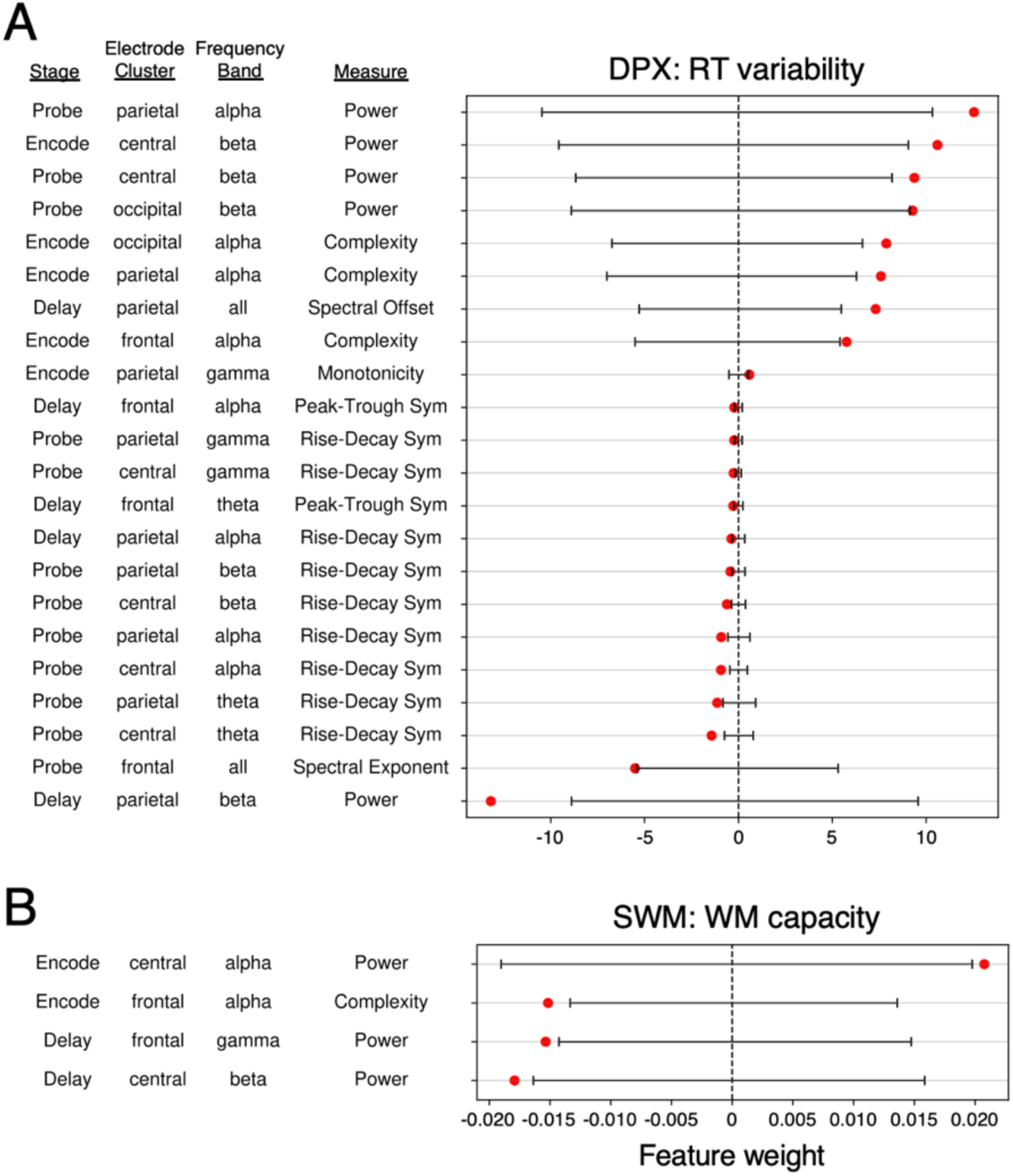
Significant features for models that significantly generalized. Significant feature coefficients (red dots) from models trained on the entire exploratory dataset (no cross-validation) for (A) DPX data predicting RT variability and (B) SWM data predicting WM capacity. Statistical significance determined as feature coefficients greater or less than the extremes of the permuted feature weight null distributions (black line spans 2.5% to 97.5% null distribution range obtained for each feature).

Across both models, power emerged as the most dominant predictor, consistent with its well-established role in the EEG literature. In the SWM model predicting WM capacity, power features overwhelmingly drove model performance, aligning with our earlier finding that the power-only model achieved significant prediction (Figure 5A, dark gray bar). Significant features were exclusively derived from encoding and delay stages. The only non-power significant feature was signal complexity, with lower neural complexity in the alpha band was associated with higher working memory capacity.

For the DPX model predicting RT variability, significant weights were concentrated on features derived from the probe stage (12 out of 22 significant predictors), consistent with the finding from our stage-specific analyses showing that a model using only probe stage EEG features achieved significant predictive performance (Figure S1). Power remained the strongest predictor but showed mixed directionality: encoding and probe power contributed positively, and delay power contributed negatively. Several complexity measures were significant, including the same alpha band complexity feature identified in the SWM model predicting WM capacity. In addition, many significant predictors reflected waveform symmetry (11 of 22 significant features).

Complexity computed from frontal alpha signal during encoding emerged as a significant predictor in both of these models. The opposite signs of coefficients arise because the behavioral outcome measures have inverse relationships: better performance is reflected by lower RT variability and higher working memory capacity. Although the feature profiles differed across tasks, the presence of an overlapping predictor suggests shared neural mechanisms supporting cognitive control across domains.

## Discussion

The present study quantified the relative contribution of several candidate EEG metrics, including spectral power as well as oscillatory structure and signal complexity, derived from working memory and resting-state recordings to predict working memory performance and psychopathology. To distinguish model optimization from model evaluation, we applied cross-validation within an exploratory dataset and reserved an independent dataset half for final validation, consistent with emerging best-practice recommendations (Poldrack et al., 2020; Scheinost et al., 2019). This framework yielded replicable brain-behavior relationships across dataset halves for cognitive outcomes of reaction time variability and working memory capacity, strengthening confidence in model robustness given that the inherent unreliability of neural and behavioral measures places an upper bound on attainable effect sizes (Marek et al., 2022). In contrast, the models predicting psychopathology – an outcome more distal to the brain activity being recorded – did not significantly generalize to the validation dataset. Oscillatory power emerged as a dominant predictor, consistent with its well-established role in cognitive function, while additional EEG measures of oscillation shape, broadband spectral structure, and signal complexity accounted for complementary variance that enhanced predictive accuracy.

### Task-evoked EEG predicts proximal working memory outcomes

The most reliable brain-behavior relationship emerged from features of EEG recorded during the DPX task, which robustly predicted RT variability on that task. RT variability is associated with sustained attention, capturing moment-to-moment fluctuations in attentional engagement that affect the stability of goal maintenance and response control (Fortenbaugh et al., 2017; Unsworth & Robison, 2018). Both unusually fast and unusually slow responses may reflect distinct lapses in attention: impulsive, premature responses often precede commission errors, whereas delayed responses frequently precede omission errors (Cheyne et al., 2009; Yamashita et al., 2021). As such, higher RT variability is generally indictive of poorer sustained attention. Elevated RT variability is frequently found in people with ADHD and related attention disorders (Machida et al., 2019; Moses et al., 2022; Tamm et al., 2012).

Among the tasks included in the present study, DPX is the least traditional working memory paradigm but is particularly well suited for indexing sustained attention. Derived from continuous performance task (CPT; Rosvold et al., 1956) paradigms, DPX emphasizes goal maintenance with context monitoring (Servan-Schreiber et al., 1996), which is a core control process that interacts closely with working memory, attention, and executive function (J. A. H. Jones et al., 2010; Lopez-Garcia et al., 2016). Unlike simpler CPT variants such as the Sustained Attention to Response Task (SART; Robertson et al., 2014), the DPX paradigm requires context-dependent inhibition: although most trials require a response, successful performance depends on maintaining contextual cues across delays and withholding responses only when they indicate a non-target pairing. In contrast, the SWM and DFR tasks involve variable memory loads that lengthen response times and engage additional cognitive resources, causing RT variability to reflect trial-to-trial fluctuations in cognitive demand (e.g., working memory load effects) rather than intrinsic attentional stability. The DPX’s consistent low-load structure minimizes these task-related fluctuations, allowing RT variability to serve as a more direct behavioral index of sustained attention, consistent with prior work linking increased reaction time variability with worse cognitive outcomes (Bielak et al., 2010; Jutten et al., 2024).

The other model that successfully generalized to the held-out dataset linked EEG features from the SWM task to individual differences in working memory capacity. Power features emerged as the most significant contributors, which is consistent with the observation that the power-only model was statistically significant within the cross-validated analyses of the exploratory dataset. In the present analysis, higher alpha and lower beta power were associated with greater working memory capacity as indicated by positive and negative model weights, respectively. This aligns with extensive evidence that modulations in oscillatory power can index core mechanisms of neural coordination supporting working memory. For example, frontal midline theta power has been associated with active maintenance and top-down cognitive control (Cavanagh & Frank, 2014; Zakrzewska & Brzezicka, 2014). Alpha power often increases during maintenance with memory load and has been interpreted as facilitating selective gating of relevant representations while suppressing interference from irrelevant inputs (Jensen et al., 2002; Klimesch et al., 2007; Manza et al., 2014; Sauseng et al., 2009). Additionally, beta power decreases have been associated with active updating and flexible cortical communication required for manipulating items in memory (Lundqvist et al., 2016; Pfurtscheller & Lopes da Silva, 1999; Spitzer & Haegens, 2017).

### EEG features supporting prediction of working memory outcomes

Among the additional EEG measures evaluated, signal complexity quantified using the Lempel-Ziv algorithm emerged as a reliable predictor of both RT variability and working memory capacity. This convergence suggests that signal complexity indexes a shared neural property supporting adaptive information processing across cognitive domains rather than a task-specific process. In both cases, lower complexity was associated with better performance (i.e., reduced RT variability and higher working memory capacity), consistent with prior findings that EEG complexity decreases under high mental load conditions (Liu et al., 1997). This pattern implies that excessive irregularity or noise in neural signals may reflect unstable coordination, whereas more ordered dynamics facilitate efficient encoding and maintenance of information. Notably, clinical populations do not exhibit a uniform direction of complexity change (Hernández et al., 2023; Yang & Tsai, 2013), suggesting that EEG signal complexity reflects neural organization supporting cognitive performance rather than a unidirectional biomarker of dysfunction. Future studies should consider how well complexity generalizes to other cognitive domains.

Oscillatory power emerged as an informative feature type across tasks. In the SWM task, power features captured meaningful predictive signal in the cross-validated exploratory analyses: both the full model and the power-only model significantly predicted working memory capacity. In contrast, for the DPX task, the power-only model was not significant in the exploratory dataset, even though many of the significant predictors of the full model were power features. The prominence of power may be understood in light of how shared variance is distributed across predictors in ridge regression. Power features were relatively independent from the more intercorrelated waveform shape and complexity measures, which likely amplified its apparent contribution within the ridge regression framework. Ridge regression distributes weight across correlated predictors while assigning larger coefficients to predictors that capture more unique variance (Cule & De Iorio, 2013; Sztepanacz & Houle, 2024). When predicting working memory capacity from SWM task data within the exploratory dataset, we found that the model excluding power features yielded numerically higher performance than the omnibus model including power. These results suggest that additional signal processing measures captured behaviorally relevant neural structure that was partially overlapping with yet complementary to information contained in power.

Oscillatory power was normalized relative to a pre-stimulus baseline to align with established conventions for quantifying task-related modulation of spectral amplitude (M. X. Cohen, 2014) and to reduce the influence of scaling factors such as anatomical and recording variability that is unrelated to neural computation (Buzsáki et al., 2012). In contrast, the additional signal processing measures of waveform shape metrics, broadband spectral characteristics, and complexity are not dependent on absolute signal amplitude but instead reflect scale-invariant or structural properties of neural activity. Because these metrics are defined in terms of ratios, temporal relationships, or signal pattern structure rather than absolute magnitude, they do not require baseline normalization in the same way as oscillatory power. As a result, baseline normalization may have further reduced shared variance between power and the other measures, increasing its apparent statistical independence within the regularized modeling framework.

### Task-evoked EEG prediction of psychopathology did not generalize

Models predicting psychopathology did not generalize to the validation dataset. Nonetheless, within the exploratory dataset, psychopathology scores were significantly predicted by EEG features derived from eyes-open rest, though these effects were weaker than those observed for task-based models. Notably, eyes-open rest involved greater cortical engagement and processing of visual input compared to the lower-arousal state captured during eyes-closed rest (Barry et al., 2007). While this result should be interpreted cautiously, it raises the possibility that psychopathology may relate more strongly to stable, trait-like neural configurations captured at rest than to transient, task-evoked activity. Indeed, resting-state EEG has long been considered a window into baseline network organization and excitation-inhibition balance, both of which are thought to be altered in neuropsychiatric conditions (Voytek & Knight, 2015; Waschke et al., 2021).

### Future directions

Several limitations and avenues for future research should be noted. Somewhat surprisingly, we were unable to reliably predict task accuracy from EEG data collected during the very performance of each task. In contrast to studies showing that neural activity can distinguish correct from incorrect response at the trial level (Johannesen et al., 2016), the present analyses focused on between-subject differences in overall accuracy, suggesting that task-evoked neural engagement may not consistently track individual differences in performance. Additionally, it is notable that EEG features derived from the SWM task predicted working memory capacity despite our capacity composite score – derived from standardized neuropsychological tests – incorporating both spatial and non-spatial behavioral measures. This result suggests that the SWM task engaged domain-general working memory processes that capture capacity beyond spatial representations alone. More broadly, the three working memory tasks evaluated in the present study (DFR, SWM, and DPX) showed largely dissociable relationships with behavioral outcomes, suggesting that they index partially distinct cognitive phenomena rather than a single, unitary working memory construct. This dissociation extended to individual task stages, which did not consistently relate to all outcome variables. This is consistent with our approach of considering working memory as the coordination of functionally distinct sub-processes. Future research could exploit these task- and stage-specific differences to clarify how distinct working memory operations underpin variability in cognitive performance and clinical outcomes. In addition, future studies should compare the predictive value of the EEG features as examined here with functional connectivity measures, which have shown promise for predicting working memory performance (Pashkov & Dakhtin, 2025). Finally, despite the clinical heterogeneity of our sample, many participants exhibited floor-level scores on psychopathology factors (indicting minimal symptom burden), limiting the variance available for prediction. Future studies aimed at discovering EEG-based biomarkers of psychopathology would benefit from sampling strategies that more evenly span the full range of symptom severity as well as longitudinal designs that track symptom emergence or change over time.

### Conclusion

Overall, we found that integrating multiple oscillatory and non-oscillatory EEG features from data measured during working memory task performance yielded reliable prediction of two cognitive outcomes: working memory capacity and response time variability. However, models that showed promise for predicting psychopathology in our exploratory dataset did not generalize when applied to our validation dataset. These findings demonstrate that incorporating diverse signal properties alongside traditional oscillatory power measures captures complementary aspects of neural dynamics that enhance our ability to explain individual differences in behavior. Had we relied solely on cross-validated performance within the exploratory dataset, we would have concluded that multiple outcomes – including psychopathology – were robustly predictable. By reserving half of the data as a fully held-out validation dataset until feature selection and model specification were finalized, we were able to more rigorously assess model reliability and reveal limitations in generalizability that would otherwise have gone undetected. This strengthens confidence in the models that did generalize and highlights the importance of strict validation procedures in neuroimaging-based prediction research.

## Acknowledgments

This work was supported by funding from the National Institute of Mental Health to R.M.B. (R01MH101478), to A.L. and S.K.L (R01MH116268), and to A.L. (R01MH128475). F.P. was supported by a National Science Foundation Graduate Research Fellowship (DGE-2034835 and DGE-2444110) and a UCLA Dissertation Year Fellowship.

## Supplementary material

**Supplemental Figure S1:**
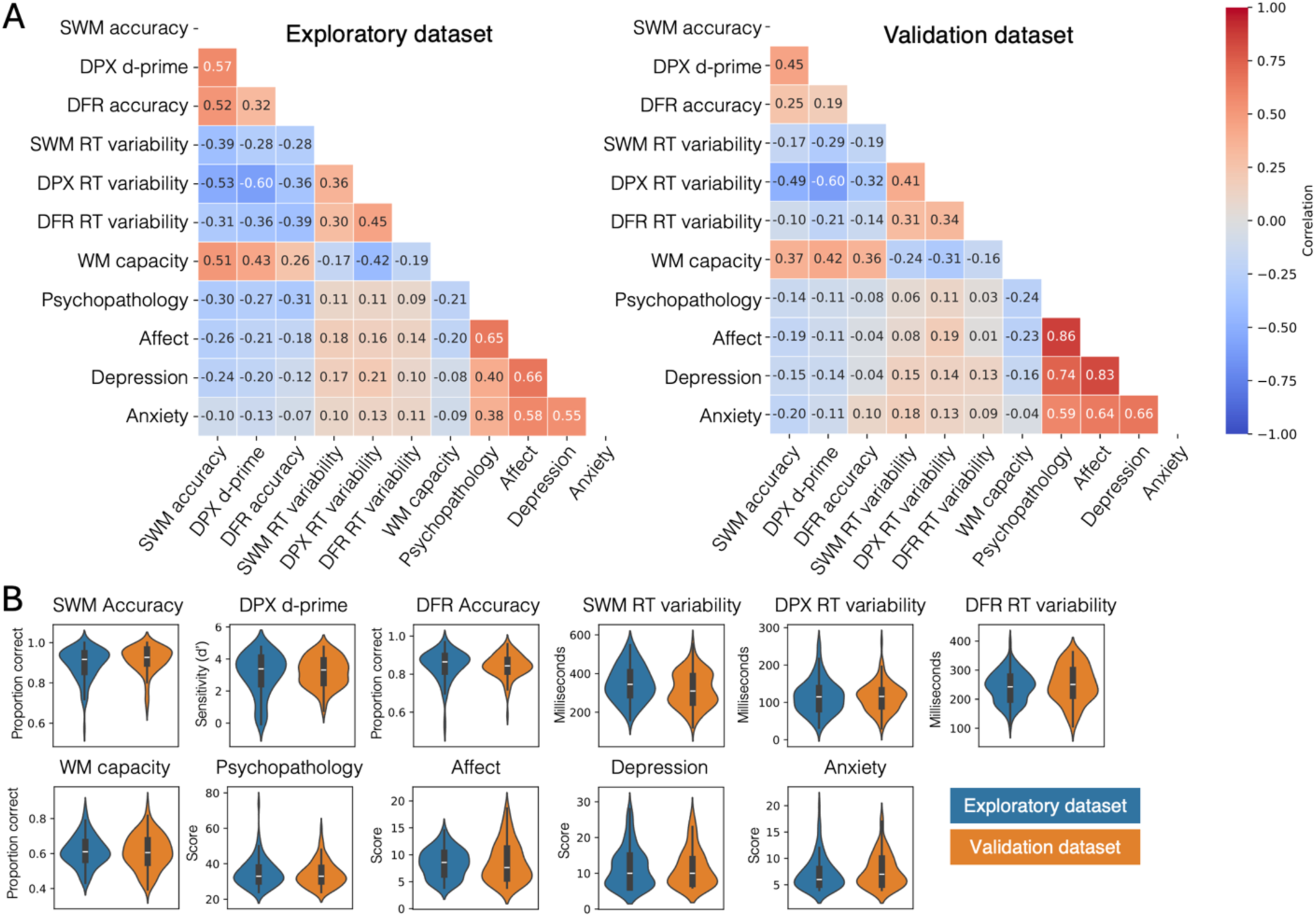
Relationship among behavioral outcome measures and comparison across datasets. (A) Heatmaps showing Pearson correlations between behavioral outcome measures within the exploratory (left) and validation (right) datasets. Each cell reflects the correlation between a pair of outcome variables. (B) Violin plots showing the distribution of each outcome measure in the exploratory (blue) and validation (orange) datasets. The width of each violin reflects the probability density of the data. Inner boxplots indicate the median and interquartile range. Distributions of all measures were comparable between the exploratory and validation datasets (all FDR-corrected *p* > .05).

**Supplemental Figure S2:**
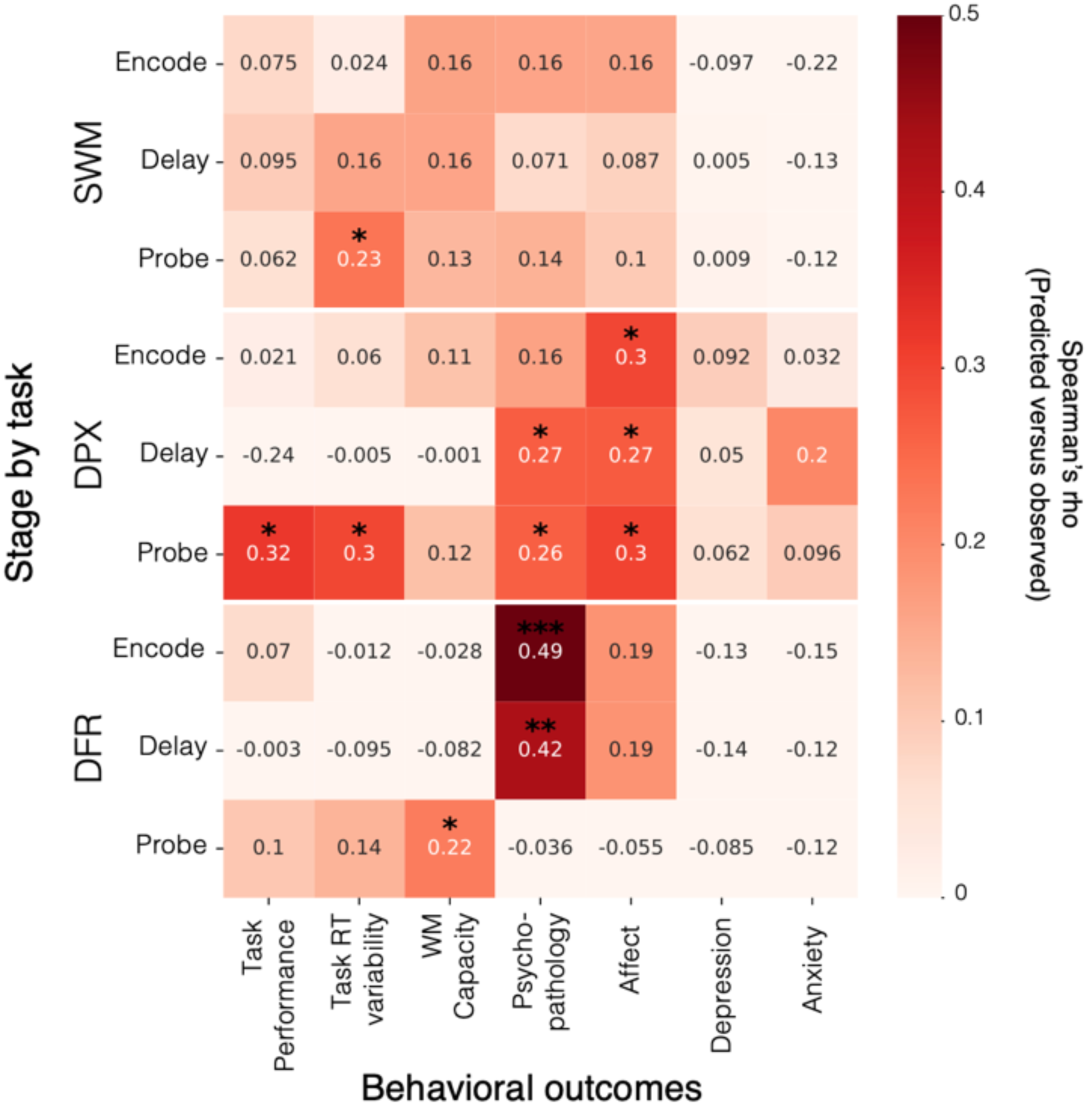
Exploratory stage-specific model performance. Heatmap of the Spearman correlation between observed and predicted values for models trained with EEG features from individual stages of each task (y-axis) for different behavioral outcome variables (x-axis). Color reflects strength of correlation that is also denoted in the center of each cell. *p<0.05; **p<0.01; ***p<0.001.

**Supplemental Figure S3:**
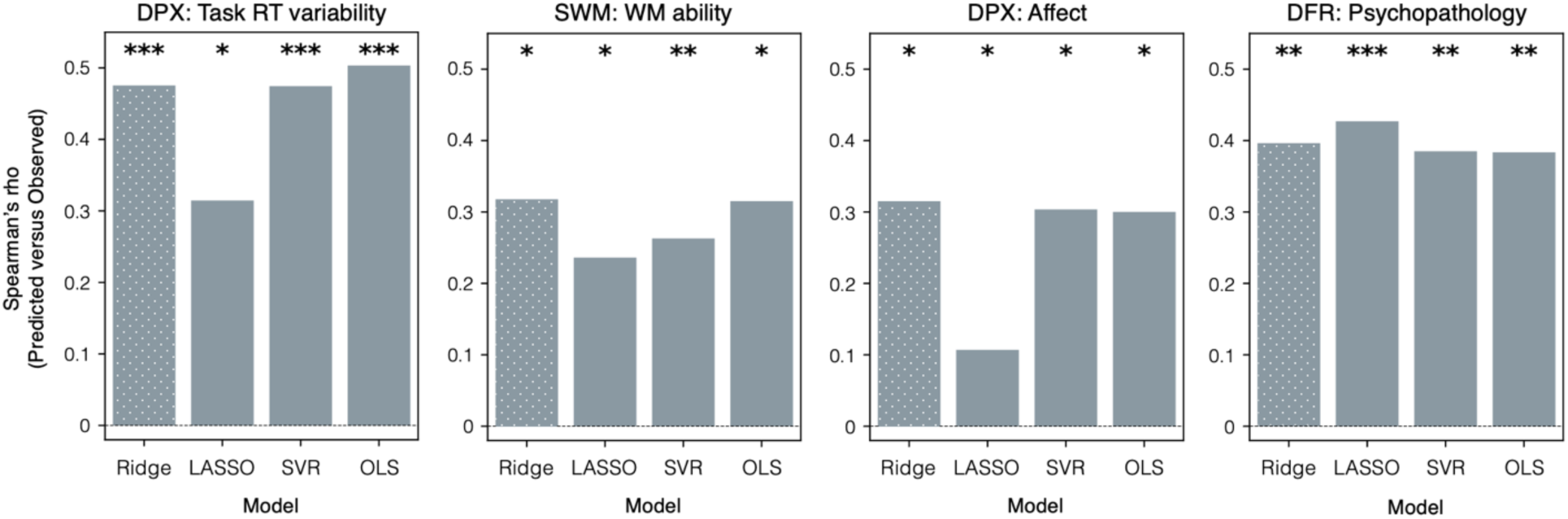
Comparison of machine learning model performanc. Performance for each main text model tested on held-out data using alternative regression approaches: ridge regression (dotted patten; model reported in main text), LASSO, support vector regression (SVR), and ordinary least squares (OLS). *p<0.05; **p<0.01; ***p<0.001.

**Supplemental Figure S4:**
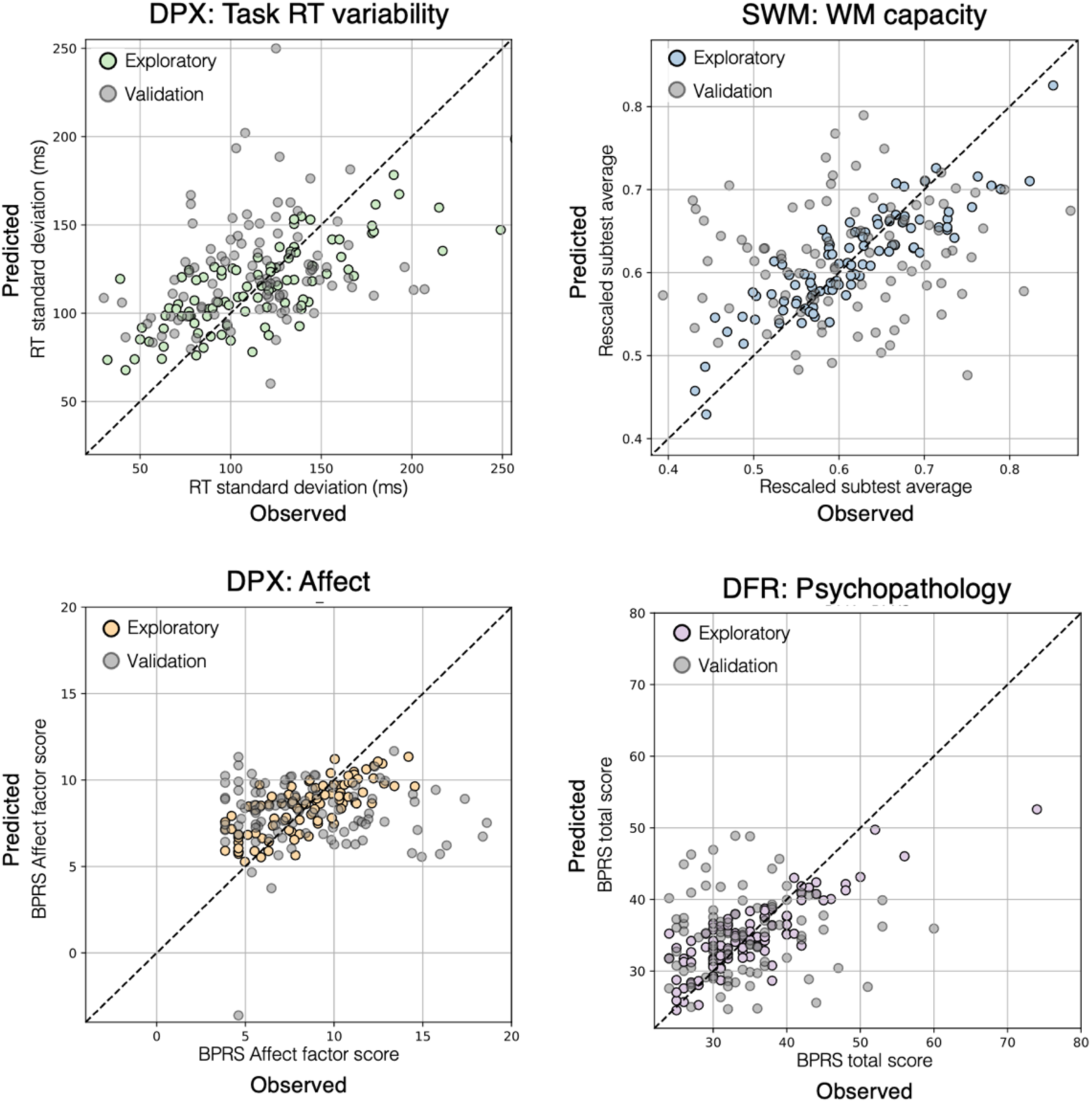
Scatterplots of validated models. Each plot shows observed versus predicted values for participants in the exploratory dataset used to train the model (colored circles) and participants in the validation dataset used to test the model (gray circles). Observed values are plotted on the x-axis and predicted values on the y-axis. Axis units correspond to the respective behavioral outcome. The diagonal dashed line indicates perfect correlation.

## References

Adam, K. C. S., Mance, I., Fukuda, K., & Vogel, E. K. (2015). The Contribution of Attentional Lapses to Individual Differences in Visual Working Memory Capacity. Journal of Cognitive Neuroscience, 27(8), 1601–1616. 10.1162/jocn_a_00811

Baddeley, A. (1992). Working Memory. Science, 255(5044), 556–559.

Barak, O., & Tsodyks, M. (2014). Working models of working memory. *Current Opinion in Neurobiology*, Theoretical and Computational Neuroscience, 25, 20–24. 10.1016/j.conb.2013.10.008

Barch, D. M., & Ceaser, A. (2012). Cognition in schizophrenia: Core psychological and neural mechanisms. Trends in Cognitive Sciences, 16(1), 27–34. 10.1016/j.tics.2011.11.015

Barry, R. J., Clarke, A. R., Johnstone, S. J., Magee, C. A., & Rushby, J. A. (2007). EEG differences between eyes-closed and eyes-open resting conditions. Clinical Neurophysiology: Official Journal of the International Federation of Clinical Neurophysiology, 118(12), 2765–2773. 10.1016/j.clinph.2007.07.028

Bender, A., Zhao, C., Vogel, E., Awh, E., & Voytek, B. (2025). Differential representations of spatial location by aperiodic and alpha oscillatory activity in working memory. Proceedings of the National Academy of Sciences, 122(30), e2506418122. 10.1073/pnas.2506418122

Bielak, A. A. M., Hultsch, D. F., Strauss, E., MacDonald, S. W. S., & Hunter, M. A. (2010). Intraindividual variability in reaction time predicts cognitive outcomes 5 years later. Neuropsychology, 24(6), 731–741. 10.1037/a0019802

Bilder, R. M., Howe, A. G., & Sabb, F. w. (2013). Multilevel models from biology to psychology: Mission impossible? Journal of Abnormal Psychology, 122(3), 917–927. 10.1037/a0032263

Bosl, W., Tierney, A., Tager-Flusberg, H., & Nelson, C. (2011). EEG complexity as a biomarker for autism spectrum disorder risk. BMC Medicine, 9(1), 18. 10.1186/1741-7015-9-18

Buzsáki, G., Anastassiou, C. A., & Koch, C. (2012). The origin of extracellular fields and currents—EEG, ECoG, LFP and spikes. Nature Reviews Neuroscience, 13(6), 407–420. 10.1038/nrn3241

Cavanagh, J. F., & Frank, M. J. (2014). Frontal theta as a mechanism for cognitive control. Trends in Cognitive Sciences, 18(8), 414–421. 10.1016/j.tics.2014.04.012

Cella, D., Riley, W., Stone, A., Rothrock, N., Reeve, B., Yount, S., Amtmann, D., Bode, R., Buysse, D., Choi, S., Cook, K., DeVellis, R., DeWalt, D., Fries, J. F., Gershon, R., Hahn, E. A., Lai, J.-S., Pilkonis, P., Revicki, D., … Hays, R. (2010). The Patient-Reported Outcomes Measurement Information System (PROMIS) developed and tested its first wave of adult self-reported health outcome item banks: 2005–2008. Journal of Clinical Epidemiology, 63(11), 1179–1194. 10.1016/j.jclinepi.2010.04.011

Cheyne, J., Solman, G. J. F., Carriere, J. S. A., & Smilek, D. (2009). Anatomy of an error: A bidirectional state model of task engagement/disengagement and attention-related errors. Cognition, 111(1), 98–113. 10.1016/j.cognition.2008.12.009

Chun, M. M. (2011). Visual working memory as visual attention sustained internally over time. *Neuropsychologia*, Attention and Short-Term Memory, 49(6), 1407–1409. 10.1016/j.neuropsychologia.2011.01.029

Cohen, J. D., Barch, D. M., Carter, C., & Servan-Schreiber, D. (1999). Context-processing deficits in schizophrenia: Converging evidence from three theoretically motivated cognitive tasks. Journal of Abnormal Psychology, 108(1), 120–133. 10.1037//0021-843x.108.1.120

Cohen, M. X. (2014). Analyzing Neural Time Series Data: Theory and Practice. MIT Press.

Cole, S. R., & Voytek, B. (2017). Brain Oscillations and the Importance of Waveform Shape. Trends in Cognitive Sciences, 21(2), 137–149. 10.1016/j.tics.2016.12.008

Cole, S. R., & Voytek, B. (2019). Cycle-by-cycle analysis of neural oscillations. Journal of Neurophysiology, 122(2), 849–861. 10.1152/jn.00273.2019

Cule, E., & De Iorio, M. (2013). Ridge regression in prediction problems: Automatic choice of the ridge parameter. Genetic Epidemiology, 37(7), 704–714. 10.1002/gepi.21750

Dazzi, F., Shafer, A., & Lauriola, M. (2016). Meta-analysis of the Brief Psychiatric Rating Scale – Expanded (BPRS-E) structure and arguments for a new version. Journal of Psychiatric Research, 81, 140–151. 10.1016/j.jpsychires.2016.07.001

D’Esposito, M., & Postle, B. R. (2015). The Cognitive Neuroscience of Working Memory. Annual Review of Psychology, 66(Volume 66, 2015), 115–142. 10.1146/annurev-psych-010814-015031

Donoghue, T., Haller, M., Peterson, E. J., Varma, P., Sebastian, P., Gao, R., Noto, T., Lara, A. H., Wallis, J. D., Knight, R. T., Shestyuk, A., & Voytek, B. (2020). Parameterizing neural power spectra into periodic and aperiodic components. Nature Neuroscience, 23(12), 1655–1665. 10.1038/s41593-020-00744-x

Eriksson, J., Vogel, E. K., Lansner, A., Bergström, F., & Nyberg, L. (2015). Neurocognitive Architecture of Working Memory. Neuron, 88(1), 33–46. 10.1016/j.neuron.2015.09.020

Esterman, M., Noonan, S. K., Rosenberg, M., & DeGutis, J. (2013). In the Zone or Zoning Out? Tracking Behavioral and Neural Fluctuations During Sustained Attention. Cerebral Cortex, 23(11), 2712–2723. 10.1093/cercor/bhs261

Fortenbaugh, F. C., DeGutis, J., & Esterman, M. (2017). Recent theoretical, neural, and clinical advances in sustained attention research. Annals of the New York Academy of Sciences, 1396(1), 70–91. 10.1111/nyas.13318

Frelih, T., Matkovič, A., Mlinarič, T., Bon, J., & Repovš, G. (2025). Modulation of aperiodic EEG activity provides sensitive index of cognitive state changes during working memory task. eLife, 13, RP101071. 10.7554/eLife.101071

Gao, R., Peterson, E. J., & Voytek, B. (2017). Inferring synaptic excitation/inhibition balance from field potentials. NeuroImage, 158, 70–78. 10.1016/j.neuroimage.2017.06.078

Hernández, R. M., Ponce-Meza, J. C., Saavedra-López, M. Á., Campos Ugaz, W. A., Chanduvi, R. M., & Monteza, W. C. (2023). Brain Complexity and Psychiatric Disorders. Iranian Journal of Psychiatry, 18(4), 493–502. 10.18502/ijps.v18i4.13637

Hoerl, A. E., & Kennard, R. W. (1970). Ridge Regression: Biased Estimation for Nonorthogonal Problems.

Hofmann, A. B., Schmid, H. M., Jabat, M., Brackmann, N., Noboa, V., Bobes, J., Garcia-Portilla, M. P., Seifritz, E., Vetter, S., & Egger, S. T. (2022). Utility and validity of the Brief Psychiatric Rating Scale (BPRS) as a transdiagnostic scale. Psychiatry Research, 314, 114659. 10.1016/j.psychres.2022.114659

Hosseini, M., Powell, M., Collins, J., Callahan-Flintoft, C., Jones, W., Bowman, H., & Wyble, B. (2020). I tried a bunch of things: The dangers of unexpected overfitting in classification of brain data. Neuroscience & Biobehavioral Reviews, 119, 456–467. 10.1016/j.neubiorev.2020.09.036

Jensen, O., Gelfand, J., Kounios, J., & Lisman, J. E. (2002). Oscillations in the Alpha Band (9–12 Hz) Increase with Memory Load during Retention in a Short-term Memory Task. Cerebral Cortex, 12(8), 877–882. 10.1093/cercor/12.8.877

Johannesen, J. K., Bi, J., Jiang, R., Kenney, J. G., & Chen, C.-M. A. (2016). Machine learning identification of EEG features predicting working memory performance in schizophrenia and healthy adults. Neuropsychiatric Electrophysiology, 2(1), 3. 10.1186/s40810-016-0017-0

Jones, J. A. H., Sponheim, S. R., & MacDonald, A. W. (2010). The dot pattern expectancy task: Reliability and replication of deficits in schizophrenia. Psychological Assessment, 22(1), 131–141. 10.1037/a0017828

Jones, S. (2016). When brain rhythms aren’t “rhythmic”: Implication for their mechanisms and meaning. Current Opinion in Neurobiology, 40, 72–80. 10.1016/j.conb.2016.06.010

Jutten, R. J., Amariglio, R. E., Maruff, P., Properzi, M. J., Rentz, D. M., Johnson, K. A., Sperling, R. A., & Papp, K. V. (2024). Increased intraindividual variability in reaction time performance is associated with emerging cognitive decline in cognitively unimpaired adults. Neuropsychology, 38(2), 184–197. 10.1037/neu0000928

Klimesch, W., Sauseng, P., & Hanslmayr, S. (2007). EEG alpha oscillations: The inhibition-timing hypothesis. Brain Research Reviews, 53(1), 63–88. 10.1016/j.brainresrev.2006.06.003

Klonowski, W. (2009). Everything you wanted to ask about EEG but were afraid to get the right answer. Nonlinear Biomedical Physics, 3, 2. 10.1186/1753-4631-3-2

Kohonen, T. (2012). Associative Memory: A System-Theoretical Approach. Springer Science & Business Media.

Lempel, A., & Ziv, J. (1976). On the Complexity of Finite Sequences. IEEE Transactions on Information Theory, 22(1), 75–81. IEEE Transactions on Information Theory. 10.1109/TIT.1976.1055501

Lenartowicz, A., Truong, H., Enriquez, K. D., Webster, J., Pochon, J.-B., Rissman, J., Bearden, C. E., Loo, S. K., & Bilder, R. M. (2021). Neurocognitive subprocesses of working memory performance. *Cognitive, Affective*, & Behavioral Neuroscience, 21(6), 1130–1152. 10.3758/s13415-021-00924-7

Lenartowicz, A., Truong, H., Salgari, G. C., Bilder, R. M., McGough, J., McCracken, J. T., & Loo, S. K. (2019). Alpha modulation during working memory encoding predicts neurocognitive impairment in ADHD. Journal of Child Psychology and Psychiatry, 60(8), 917–926. 10.1111/jcpp.13042

Liu, J., He, T., Zheng, C., & Huang, Y. (1997). [Measuring EEG complexity for studying the state of mental load]. Sheng Wu Yi Xue Gong Cheng Xue Za Zhi = Journal of Biomedical Engineering = Shengwu Yixue Gongchengxue Zazhi, 14(1), 33–37.

Lopez-Garcia, P., Lesh, T. A., Salo, T., Barch, D. M., MacDonald, A. W., Gold, J. M., Ragland, J. D., Strauss, M., Silverstein, S. M., & Carter, C. S. (2016). The neural circuitry supporting goal maintenance during cognitive control: A comparison of expectancy AX-CPT and dot probe expectancy paradigms. *Cognitive, Affective*, & Behavioral Neuroscience, 16(1), 164–175. 10.3758/s13415-015-0384-1

Lundqvist, M., Rose, J., Herman, P., Brincat, S. L., Buschman, T. J., & Miller, E. K. (2016). Gamma and Beta Bursts Underlie Working Memory. Neuron, 90(1), 152–164. 10.1016/j.neuron.2016.02.028

Machida, K., Murias, M., & Johnson, K. A. (2019). Electrophysiological Correlates of Response Time Variability During a Sustained Attention Task. Frontiers in Human Neuroscience, 13. 10.3389/fnhum.2019.00363

Manza, P., Hau, C. L. V., & Leung, H.-C. (2014). Alpha Power Gates Relevant Information during Working Memory Updating. Journal of Neuroscience, 34(17), 5998–6002. 10.1523/JNEUROSCI.4641-13.2014

Marek, S., Tervo-Clemmens, B., Calabro, F. J., Montez, D. F., Kay, B. P., Hatoum, A. S., Donohue, M. R., Foran, W., Miller, R. L., Hendrickson, T. J., Malone, S. M., Kandala, S., Feczko, E., Miranda-Dominguez, O., Graham, A. M., Earl, E. A., Perrone, A. J., Cordova, M., Doyle, O., … Dosenbach, N. U. F. (2022). Reproducible brain-wide association studies require thousands of individuals. Nature, 603(7902), 654–660. 10.1038/s41586-022-04492-9

Martinussen, R., Hayden, J., Hogg-johnson, S., & Tannock, R. (2005). A Meta-Analysis of Working Memory Impairments in Children With Attention-Deficit/Hyperactivity Disorder. Journal of the American Academy of Child & Adolescent Psychiatry, 44(4), 377–384. 10.1097/01.chi.0000153228.72591.73

McCabe, D. P., Roediger III, H. L., McDaniel, M. A., Balota, D. A., & Hambrick, D. Z. (2010). The relationship between working memory capacity and executive functioning: Evidence for a common executive attention construct. Neuropsychology, 24(2), 222–243. 10.1037/a0017619

McKeon, S. D., Perica, M. I., Parr, A. C., Calabro, F. J., Foran, W., Hetherington, H., Moon, C.-H., & Luna, B. (2024). Aperiodic EEG and 7T MRSI evidence for maturation of E/I balance supporting the development of working memory through adolescence. Developmental Cognitive Neuroscience, 66, 101373. 10.1016/j.dcn.2024.101373

McTeague, L. M., Goodkind, M. S., & Etkin, A. (2016). Transdiagnostic impairment of cognitive control in mental illness. Journal of Psychiatric Research, 83, 37–46. 10.1016/j.jpsychires.2016.08.001

Miller, E. K., & Cohen, J. D. (2001). An Integrative Theory of Prefrontal Cortex Function. Annual Review of Neuroscience, 24(Volume 24, 2001), 167–202. 10.1146/annurev.neuro.24.1.167

Moses, M., Tiego, J., Demontis, D., Bragi Walters, G., Stefansson, H., Stefansson, K., Børglum, A. D., Arnatkeviciute, A., & Bellgrove, M. A. (2022). Working memory and reaction time variability mediate the relationship between polygenic risk and ADHD traits in a general population sample. Molecular Psychiatry, 27(12), 5028–5037. 10.1038/s41380-022-01775-5

Myers, N. E., Stokes, M. G., & Nobre, A. C. (2017). Prioritizing Information during Working Memory: Beyond Sustained Internal Attention. Trends in Cognitive Sciences, 21(6), 449–461. 10.1016/j.tics.2017.03.010

Newson, J. J., & Thiagarajan, T. C. (2018). EEG Frequency Bands in Psychiatric Disorders: A Review of Resting State Studies. Frontiers in Human Neuroscience, 12, 521. 10.3389/fnhum.2018.00521

Overall, J. E., & Gorham, D. R. (1962). *Brief Psychiatric Rating Scale* [Dataset]. 10.1037/t01554-000

Pashkov, A., & Dakhtin, I. (2025). Direct Comparison of EEG Resting State and Task Functional Connectivity Patterns for Predicting Working Memory Performance Using Connectome-Based Predictive Modeling. Brain Connectivity, 15(4), 175–187. 10.1089/brain.2024.0059

Pavlov, Y. G., & Kotchoubey, B. (2022). Oscillatory brain activity and maintenance of verbal and visual working memory: A systematic review. Psychophysiology, 59(5), e13735. 10.1111/psyp.13735

Pedregosa, F., Varoquaux, G., Gramfort, A., Michel, V., Thirion, B., Grisel, O., Blondel, M., Müller, A., Nothman, J., Louppe, G., Prettenhofer, P., Weiss, R., Dubourg, V., Vanderplas, J., Passos, A., Cournapeau, D., Brucher, M., Perrot, M., & Duchesnay, É. (2018). *Scikit-learn: Machine Learning in Python* (arXiv:1201.0490). arXiv. 10.48550/arXiv.1201.0490

Pfurtscheller, G., & Lopes da Silva, F. H. (1999). Event-related EEG/MEG synchronization and desynchronization: Basic principles. Clinical Neurophysiology: Official Journal of the International Federation of Clinical Neurophysiology, 110(11), 1842–1857. 10.1016/s1388-2457(99)00141-8

Poldrack, R. A., Huckins, G., & Varoquaux, G. (2020). Establishment of Best Practices for Evidence for Prediction: A Review. JAMA Psychiatry, 77(5), 534–540. 10.1001/jamapsychiatry.2019.3671

Psychology Software Tools, Inc. (2016). E-Prime 3.0. https://support.pstnet.com

Robertson, I. H., Manly, T., Andrade, J., Baddeley, B. T., & Yiend, J. (2014). Sustained Attention to Response Task [Dataset]. 10.1037/t28308-000

Rosvold, H. E., Mirsky, A. F., Sarason, I., Bransome Jr., E. D., & Beck, L. H. (1956). A continuous performance test of brain damage. Journal of Consulting Psychology, 20(5), 343–350. 10.1037/h0043220

Roux, F., & Uhlhaas, P. J. (2014). Working memory and neural oscillations: Alpha–gamma versus theta–gamma codes for distinct WM information? Trends in Cognitive Sciences, 18(1), 16–25. 10.1016/j.tics.2013.10.010

Sauseng, P., Klimesch, W., Heise, K. F., Gruber, W. R., Holz, E., Karim, A. A., Glennon, M., Gerloff, C., Birbaumer, N., & Hummel, F. C. (2009). Brain Oscillatory Substrates of Visual Short-Term Memory Capacity. Current Biology, 19(21), 1846–1852. 10.1016/j.cub.2009.08.062

Scheinost, D., Noble, S., Horien, C., Greene, A. S., Lake, E. M., Salehi, M., Gao, S., Shen, X., O’Connor, D., Barron, D. S., Yip, S. W., Rosenberg, M. D., & Constable, R. T. (2019). Ten simple rules for predictive modeling of individual differences in neuroimaging. NeuroImage, 193, 35–45. 10.1016/j.neuroimage.2019.02.057

Servan-Schreiber, D., Cohen, J. D., & Steingard, S. (1996). Schizophrenic Deficits in the Processing of Context: A Test of a Theoretical Model. Archives of General Psychiatry, 53(12), 1105–1112. 10.1001/archpsyc.1996.01830120037008

Snyder, H. R. (2013). Major depressive disorder is associated with broad impairments on neuropsychological measures of executive function: A meta-analysis and review. Psychological Bulletin, 139(1), 81–132. 10.1037/a0028727

Spitzer, B., & Haegens, S. (2017). Beyond the Status Quo: A Role for Beta Oscillations in Endogenous Content (Re)Activation. eNeuro, 4(4), ENEURO.0170-17.2017. 10.1523/ENEURO.0170-17.2017

Sternberg, S. (1966). High-Speed Scanning in Human Memory. Science, 153(3736), 652–654. 10.1126/science.153.3736.652

Sztepanacz, J. L., & Houle, D. (2024). Regularized regression can improve estimates of multivariate selection in the face of multicollinearity and limited data. Evolution Letters, 8(3), 361–373. 10.1093/evlett/qrad064

Tamm, L., Narad, M. E., Antonini, T. N., O’Brien, K. M., Hawk, L. W., & Epstein, J. N. (2012). Reaction Time Variability in ADHD: A Review. Neurotherapeutics, 9(3), 500–508. 10.1007/s13311-012-0138-5

Unsworth, N., & Robison, M. K. (2018). Tracking arousal state and mind wandering with pupillometry. *Cognitive, Affective*, & Behavioral Neuroscience, 18(4), 638–664. 10.3758/s13415-018-0594-4

Virtue-Griffiths, S., Fornito, A., Thompson, S. C. H., Biabani, M., Tiego, J., Thapa, T., Bailey, N. W., & Rogasch, N. C. (2025). Task-related changes in aperiodic activity are related to visual working memory capacity independent of event-related potentials and alpha oscillations. Imaging Neuroscience, 3, IMAG.a.150. 10.1162/IMAG.a.150

Voytek, B., & Knight, R. T. (2015). Dynamic Network Communication as a Unifying Neural Basis for Cognition, Development, Aging, and Disease. *Biological Psychiatry*, Cortical Oscillations for Cognitive/Circuit Dysfunction in Psychiatric Disorders, 77(12), 1089–1097. 10.1016/j.biopsych.2015.04.016

Vul, E., Harris, C., Winkielman, P., & Pashler, H. (2009). Puzzlingly High Correlations in fMRI Studies of Emotion, Personality, and Social Cognition. Perspectives on Psychological Science: A Journal of the Association for Psychological Science, 4(3), 274–290. 10.1111/j.1745-6924.2009.01125.x

Waschke, L., Donoghue, T., Fiedler, L., Smith, S., Garrett, D. D., Voytek, B., & Obleser, J. (2021). Modality-specific tracking of attention and sensory statistics in the human electrophysiological spectral exponent. eLife, 10, e70068. 10.7554/eLife.70068

Yamashita, A., Rothlein, D., Kucyi, A., Valera, E. M., Germine, L., Wilmer, J., DeGutis, J., & Esterman, M. (2021). Variable rather than extreme slow reaction times distinguish brain states during sustained attention. Scientific Reports, 11(1), 14883. 10.1038/s41598-021-94161-0

Yang, A. C., & Tsai, S.-J. (2013). Is mental illness complex? From behavior to brain. Progress in Neuro-Psychopharmacology & Biological Psychiatry, 45, 253–257. 10.1016/j.pnpbp.2012.09.015

Zakrzewska, M. Z., & Brzezicka, A. (2014). Working memory capacity as a moderator of load-related frontal midline theta variability in Sternberg task. Frontiers in Human *Neuroscience*, *8*. 10.3389/fnhum.2014.00399

